# Exposing New Taxonomic Variation with Inflammation – A Murine Model-Specific Genome Database for Gut Microbiome Researchers

**DOI:** 10.1101/2022.10.24.513540

**Authors:** Ikaia Leleiwi, Josue Rodriguez-Ramos, Michael Shaffer, Anice Sabag-Daigle, Katherine Kokkinias, Rory M Flynn, Rebecca A Daly, Linnea FM Kop, Lindsey M Solden, Brian M. M. Ahmer, Mikayla A Borton, Kelly C Wrighton

**Affiliations:** Department of Cell and Molecular Biology, The Colorado State University, Fort Collins, CO, USA; Department of Soil and Crop Sciences, The Colorado State University, Fort Collins, CO, USA; Graduate Degree Program in Ecology, The Colorado State University, Fort Collins, CO, USA; Department of Microbial Infection and Immunity, The Ohio State University, Columbus, OH, USA; Department of Microbiology, Immunology, and Pathology, The Colorado State University, Fort Collins, CO, USA; Department of Microbiology, RIBES, Radbound University, Nijmegen, The Netherlands & Department of Microbiology & Biophysics, The Ohio State University, Columbus, OH, USA

**Keywords:** CBA/J mouse, Salmonella, Metagenome, Pathogen, Pathobiome

## Abstract

**Background:** The murine CBA/J mouse model widely supports immunology and enteric pathogen research. This model has illuminated *Salmonella* interactions with the gut microbiome since pathogen proliferation does not require disruptive pretreatment of the native microbiota, nor does it become systemic, thereby representing an analog to gastroenteritis disease progression in humans. Despite the value to broad research communities, microbiota in CBA/J mice are not represented in current murine microbiome genome catalogs.

**Results:** Here we present the first microbial and viral genomic catalog of the CBA/J murine gut microbiome. Using fecal microbial communities from untreated and *Salmonella*-infected, highly inflamed mice, we performed genomic reconstruction to determine the impacts on gut microbiome membership and functional potential. From high depth whole community sequencing (~42.4 Gbps/sample), we reconstructed 2,281 bacterial and 4,129 viral draft genomes. *Salmonella* challenge significantly altered gut membership in CBA/J mice, revealing 30 genera and 98 species that were conditionally rare and unsampled in non-inflamed mice. Additionally, inflamed communities were depleted in microbial genes that modulate host anti-inflammatory pathways and enriched in genes for respiratory energy generation. Our findings suggest decreases in butyrate concentrations during Salmonella infection corresponded to reductions in the relative abundance in members of the *Alistipes*. Strain-level comparison of CBA/J microbial genomes to prominent murine gut microbiome databases identified newly sampled lineages in this resource, while comparisons to human gut microbiomes extended the host relevance of dominant CBA/J inflammation resistant strains.

**Conclusions:** This CBA/J microbiome database provides the first genomic sampling of relevant, uncultivated microorganisms within the gut from this widely used laboratory model. Using this resource, we curated a functional, strain-resolved view on how *Salmonella* remodels intact murine gut communities, advancing pathobiome understanding beyond inferences from prior amplicon-based approaches. *Salmonella-*induced inflammation suppressed *Alistipes* and other dominant members, while rarer commensals like *Lactobacillus* and *Enterococcus* endure. The rare and novel species sampled across this inflammation gradient advance the utility of this microbiome resource to benefit the broad research needs of the CBA/J scientific community, and those using murine models for understanding the impact of inflammation on the gut microbiome more generally.

## Introduction

Non-typhoidal *Salmonella* (NTS) is one of the leading causes of gastroenteritis and associated mortality worldwide, resulting in nearly 1 million cases and more than 50 thousand deaths in 2017 [1–3]. *Salmonella enterica* serovar Typhimurium (referred to hereon as *Salmonella*) is a NTS model enteric pathogen that exploits inflammation to increase its pathogenicity and fitness relative to other bacteria [4–6]. Prior research in murine models showed gut microbiota are remodeled during *Salmonella*-induced inflammation because of innate immunity, diminished resources, and altered chemical environment [6,7]. Similar inflammation-associated changes in commensal gut ecology are observed in patients with Chron’s disease, irritable bowel disease, and metabolic syndrome [4,8]. The *Salmonella* disease model may represent a human-relevant system for investigating host-pathogen-microbiota interactions in the inflamed GI tract germane to microbiome changes during other human chronic inflammatory conditions.

Earlier work showed that *Salmonella* induced inflammation created reactive oxygen and nitrogen species in the luminal environment, increasing the availability of oxygen and potentiating formation of tetrathionate and nitrate [9]. Increased concentrations of these terminal electron acceptors allowed *Salmonella* to respire and out-compete obligate fermentative commensals like members of the *Clostridia* [6,10]. Additionally, in the remodeled gut ecosystem respiring *Salmonella* benefited from unique access to non-competitive carbon sources like propionate and ethanolamine [11,12]. Prior competition studies often focused on decreases in the *Clostridia* [7,13], ignoring implications for the other members of the community, and those that can withstand *Salmonella* infection. Here we provide a holistic, strain resolved view of the changes in microbiome membership and function during *Salmonella* infection.

Previous investigations of the impacts of *Salmonella* on the gut microbiome relied on the use of pre-treated reduced diversity communities [14–16]. Mice in past studies (e.g., BALB/c or C57BL/6) required antibiotic conditioning prior to pathogen introduction, preventing investigation of *Salmonella* physiology in response to an intact microbial community [14,15,17]. As an alternative, CBA/J mice are gaining appreciation as a model to interrogate *Salmonella* pathogenicity as they support longer non-systemic *Salmonella* infection without prior antibiotic perturbation, similar to enteric disease manifestation in humans [17–20]. While *in vitro* and *in vivo* studies with reduced microbiota consortia provide an important theoretical framework, additional research is needed in the presence of native microbial communities to understand how specific inflammation-induced changes to microbiota membership and function impact *Salmonella* physiology and pathogenicity.

Currently the capability to study intact resident gut microbiota during *Salmonella*-induced inflammation is hindered because CBA/J mice are missing from available murine microbiome genomic catalogs [21–23]. Furthermore, existing murine gut genomic databases exclude inflamed mice, and mice colonized by enteric pathogens (e.g. like *Salmonella, Klebsiella*, and *Citrobacter)*, thus limiting the extension of these existing microbiome resources to pathobiome models. Despite being recognized as important contributors to human health [24], these existing curated murine gut microbial genome catalogs also lack virome sampling [25,26]. Accurate interrogation of complete microbial community functions during *Salmonella* infection requires comprehensive model-specific knowledge of gene content and community membership both in healthy and inflamed guts.

To evaluate the functional potential of microbial communities during *Salmonella*-induced inflammation, and to explore if the CBA/J inflammation model harbors unique and previously understudied microorganisms, we constructed a metagenome assembled genome catalog from healthy and *Salmonella* infected CBA/J mice. We employed high depth metagenomic sequencing and used several assembly strategies to increase the *de novo* reconstruction of viral and bacterial genomes. These efforts resulted in a comprehensive culture-independent genome resource that (i) revealed novel taxonomy unique to CBA/J and inflamed mice, (ii) included taxa with relevance to human systems and with anti-inflammatory effector potential, and (iii) showed how enteric inflammation remodels the functional profile of the gut by selecting for bacteria that encode mechanisms to withstand oxidative stress. Ultimately our findings reframe existing responses of the microbiota during *Salmonella* infection and provide new insights into specific bacteria that can withstand inflammation to maintain critical gut functions, perhaps revealing promising future probiotic targets.

## Results

### Pathogen perturbation extends the genomic sampling of the CBA/J microbiome

To examine the microbial community response to *Salmonella* colonization, 14 CBA/J mice were infected with 10^9^ CFU *Salmonella enterica* serovar Typhimurium strain 14028 with results compared to 16 uninoculated control mice sampled at the same time points (n=30 mice, Fig. 1A). Feces were collected prior to infection (Day −1) and in late stages of infection (Days 10 and 11), with 16S rRNA microbial community analyses performed at early and late time points (n=60) and lipocalin-2, an indicator of enteric inflammation, measured on late time points from select mice from each treatment group (n=12). The 60 fecal samples yielded 2,047,287 paired end 16S rRNA reads, which identified 23,022 unique amplicon sequencing variants (ASVs) from both inflamed and control treatments (Data S1). To confirm infection, we established that inoculated mice had *Salmonella* relative abundance greater than 25% on Day 11 and had significantly higher lipocalin-2 concentrations than control mice on Day 10. From these mice we selected feces at Day 11 from 3 *Salmonella* infected mice and 3 uninfected mice for deep metagenomic sequencing.

**Fig. 1.**
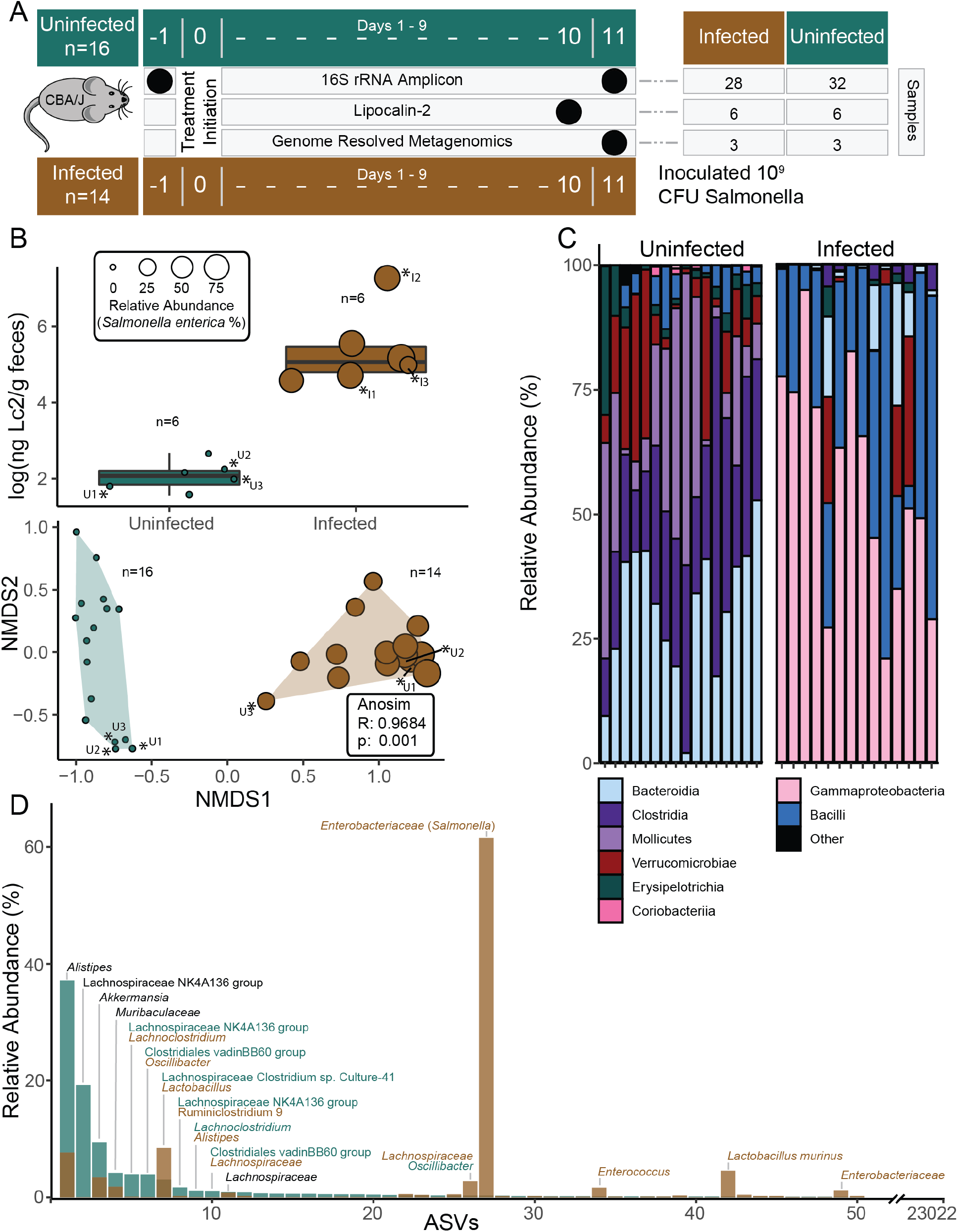
Amplicon sequencing of the CBA/J gut reveals shifts in microbiome composition with inflammation. A) Experimental design shows the number of mice in healthy (green) and infected (brown, infected with 10^9^ CFU *Salmonella*) treatments with fecal sampling times and corresponding analysis indicated by black circles. B) Boxplots show lipocalin-2 (Lc2) levels of mice in each treatment during peak infection (top), with *Salmonella* relative abundance indicated by circle size. Non-metric multidimensional scaling of Amplicon Sequence Variant (ASV) Bray-Curtis distances showing significantly distinct communities between treatments at time of sampling with points scaled to *Salmonella* relative abundance (bottom). Asterisks indicate mice used to create the CBAJ-DB. C) ASV Class distribution is depicted by stacked bar charts of healthssy and infected communities, with each bar representing a single mouse at the Day 11 timepoint. D) Rank abundance curve of mean ASV relative abundance by treatment in mice sampled to create the CBAJ-DB. Bars represent a single ASV and are colored by treatment, with ASVs ranked separately for each treatment to show the changes in rank and abundance with inflammation. Bars are labeled with taxonomic identity in text if greater than 3% mean relative abundance in either treatment, with text color denoting treatment. Black text indicates both treatments have the same ASV taxonomy within that rank.

The 16S rRNA gene findings confirmed *Salmonella* infection resulted in statistically discernable microbial communities by Day 11 following infection (Fig. 1B). A *Salmonella enterica* relative abundance increase (≥25%) was concomitant with increased inflammation evidenced by a 2.5 log-fold rise in lipocalin-2 compared to levels in uninfected mice (Fig. 1B, Data S1). The microbial community of inflamed mice statistically differed from control uninfected mice at the same time point, and pre-pathogen treated mice from both treatments (Fig. 1B, Fig. 1C, Fig. S1). Pre-infection (Day-1) mice that later became *Salmonella* inoculated, and uninoculated mice had fecal microbial communities that were not discernable from each other, indicating observed community differences by Day 11 were due to *Salmonella* infection. As others have reported [13,27], *Salmonella*-induced inflammation significantly changed gut microbial diversity, it reduced ASV richness by more than half (76.2%) and decreased Shannon’s diversity by 2.6-fold. These findings demonstrate that pathogen perturbation changes microbial membership and structure, offering a strategy for differential genomic sampling of the CBA/J gut microbiome.

These 16S rRNA analyses revealed inflamed communities were enriched in members of the *Gammaproteobacteria* and *Bacilli*, while gut communities of uninoculated mice included higher relative abundances of *Bacteroidia, Mollicutes*, and *Clostridia* (Fig. 1C). From the mice also sampled for metagenomic analysis (n=6), *Alistipes* sp. was the most dominant commensal and the most reduced during inflammation (from 37.2%), but notably still detectable (7.68%). *Salmonella enterica* Typhimurium dominated the *Gammaproteobacteria* in inflamed communities, contributing up to a mean relative abundance of 94% in infected samples. Certain low abundant members of the CBA/J microbiome significantly increased in relative abundance following pathogen treatment, including some members of *Lactobacillus, Enterococcus*, and *Lachnospiraceae* (Fig. 1D). Control mouse communities are consistent with findings from prior work showing uninfected CBA/J mouse gut community membership dominated by *Bacteroidetes* and *Firmicutes*, especially *Clostridia* of various *Lachnospiraceae* and *Ruminococcaceae* families [27,28]. These 16S rRNA gene analyses revealed abundant members in both healthy and inflamed CBA/J gut microbiomes that represented microorganismal genome ‘targets’ for our database.

### Microbial genomic reconstruction from CBA/J mice recovers relevant members sampled in amplicon surveys

To thoroughly catalog the CBA/J gut microbiota high sequencing depth was required to sequence through *Salmonella* dominance (25.8 – 94.2% by amplicon analyses) and recover some of the first genomes from rare, but persistent co-occurring members of the pathogen inflamed gut. We obtained 254.2 Gbps of metagenomic sequencing data (Data S2) from 6 representative mice (inflamed n=3, uninfected n=3, Fig. 1A), 77-fold more sequencing/sample than is commonly done in murine catalogs (Fig. S2A). Additionally, we used iterative, targeted assembly approaches (single, co-assembly, subtractive assembly) as well as two different assemblers to attempt to enhance genome quality and recovery, especially from less dominant members (Fig. 2A, Fig. S2B). Subtractive and co-assembly methods derived 259 additional metagenomically assembled genomes (MAGs) beyond those from single sample assemblies, with the distribution of MAGs from each assembler reported (Fig. 2A). In total, we recovered 2,281 MAGs. Quality assessment revealed 504 MAGs to be either medium or high-quality (MQHQ) with sufficiently low contamination to be included in further analyses. These quality genomes contained 156,921 uniquely called predicted genes [29,30] (Fig. 2, Data S3).

**Fig. 2.**
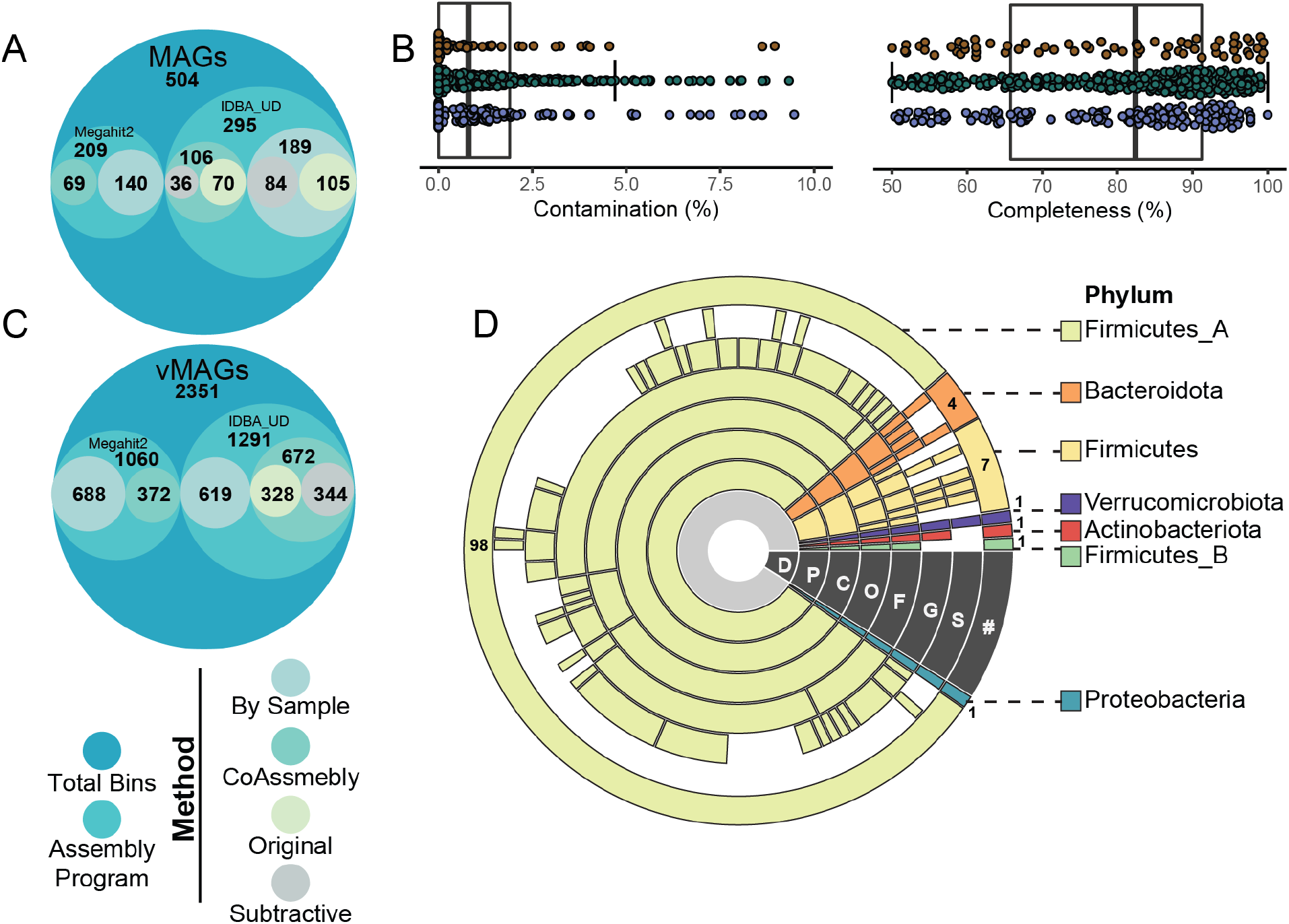
CBA/J mouse database (CBAJ-DB) genomic methods and composition. A) Circle plot shows the number of medium and high quality (MQHQ) metagenome assembled genomes (MAGs) reconstructed from CBA/J mouse gut metagenomes and corresponding assembly method and assembly software. B) Completeness and contamination of all MQHQ MAGs colored by sample treatment origin (brown = infected; green = uninfected; purple = co-assembly) are shown by box plots, with bold horizontal lines indicating median across all MQHQ MAGs. C) Circle plot shows the number of viral metagenome assembled genomes (vMAGs) reconstructed from CBA/J mouse gut metagenomes and corresponding assembly method and assembly software. D) Dereplicated MQHQ (dMQHQ) MAG phylum distribution is shown by sequential colored rings listed from least specific (Domain, D) to most specific (Species, S) moving outwards from the plot center. Gaps at each taxonomic level represent MAGs that are previously undescribed. The outer ring is labeled with the number of MAGs from the dMQHQ CBAJ-DB database within each Phylum.

Dereplication of our metagenome assembled genomes (99% identity) resulted in 113 medium and high-quality MAGs (dMQHQ) from both treatments. These MAGs were assigned to 7 Phyla – *Actinobacteriota* (1), *Bacteroidota* (4) *Firmicutes* (7), *Firmicutes*_A (98), *Firmicutes*_B (1), *Proteobacteria* (1), and *Verrucomicrobiota* (1) (Fig. 2D, Data S2). Nearly a third (30 of the 113) of the dereplicated MAGs were assigned to 30 genera and 98 to species that were only recognized by alphanumeric numbering in GTDB-Tk, hinting that novelty sampled here may be undescribed not only in murine but larger MAG collections. Reflecting the richness of these samples, the majority of MAGs originated from uninfected mice (59%) and their co-assemblies (35%) while 13% came from inflamed mice. Specifically, *Enterococcus_D gallinarum, Erysipelatoclostridium cocleatum, Kineothrix sp000403275, and Lactobacillus_B animalis* MAGs were uniquely recovered from inflamed mice, consistent with their 16S rRNA membership (Fig. 1D).

This finding indicates how perturbation can aid in the sampling of genomes from conditionally rare members. Expanding this resource beyond solely bacterial genomes, we also reconstructed viral genomes from our CBA/J assemblies, recovering 4,129 viral metagenome assembled genomes (vMAGs). Of these, 2,351 vMAGs were ≥10kb which were then dereplicated into 609 viral genomes (Fig. 2C, Fig. 4D, Data S4).

**Fig. 4.**
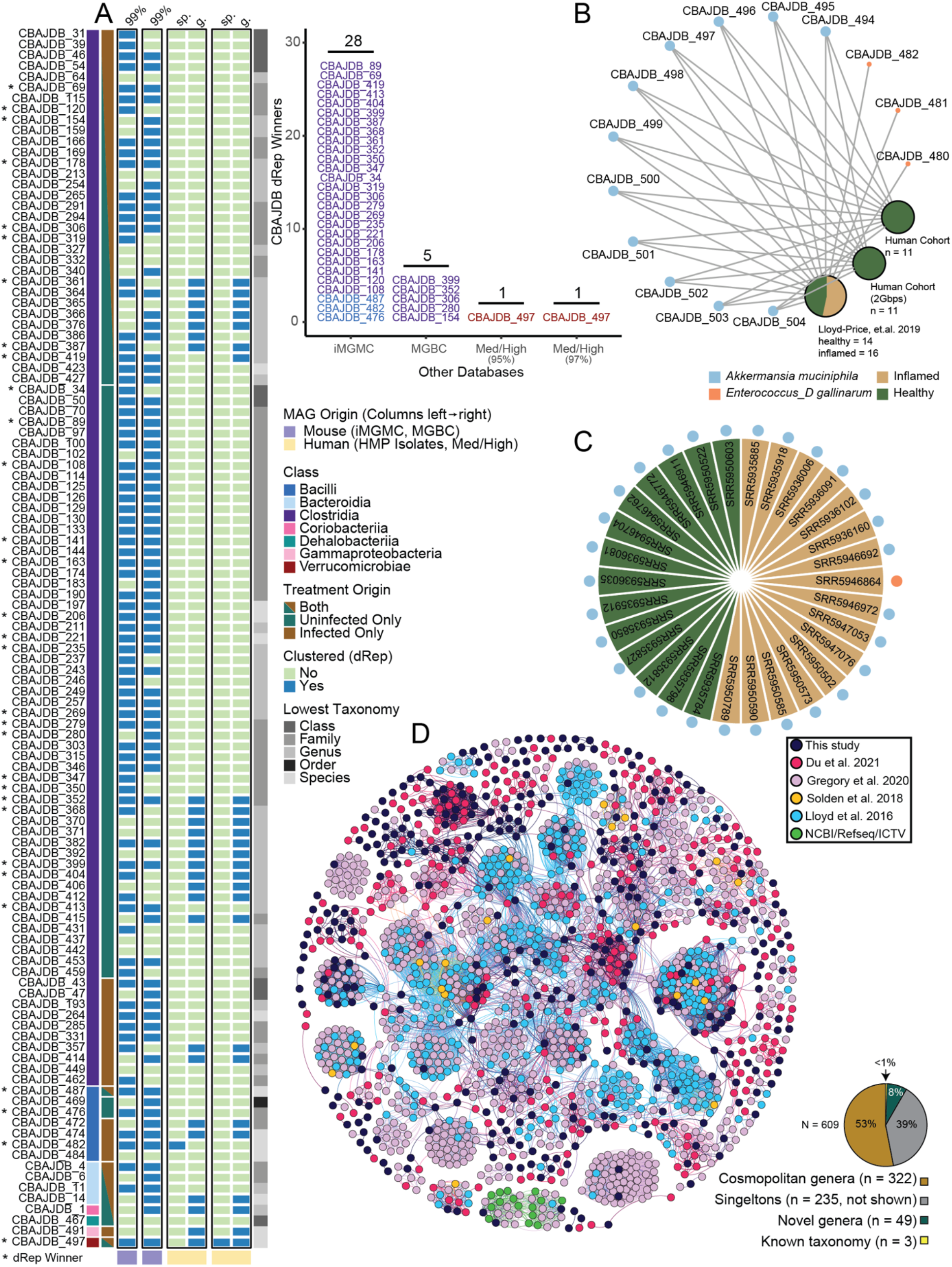
CBA/J database (CBAJ-DB) genomes link to other murine and human studies. A) Medium and high quality (MQHQ) metagenome assembled genomes (MAGs) that clustered with genomes from either murine databases (light purple columns) or the HMP/Human cohort (yellow columns). CBAJ-DB MAGs that were the highest quality genome in each cluster are marked with an asterisk and are displayed in the stacked bar chart grouped by database. MAGs are grouped by class (first bar annotation) and treatment origin (second bar annotation), with the lowest assigned taxonomy indicated in gray scale (right bar). For each database in black outline, blue cells indicate CBAJ-DB MAGs that clustered, while green cells show no clustering. Accompanying bar chart shows the number of CBAJ-DB MAGs with higher quality scores corresponding genomes in other databases. B) CBAJ-DB MAGs that recruited human reads are shown by blue (*Akkermansia muciniphila*) and orange nodes (*Enterococcus_D gallinarum*), with size indicating number of CBAJ-DB MAGs. MAG nodes are linked to databases shown as pie charts, where green (healthy human) and tan (inflamed human) indicate sequencing origin of mapped reads and node size indicates sample number. C) SRA accession ID’s of samples from the Lloyd-Price cohort that mapped to CBAJ-DB MAGs *Akkermansia muciniphila* (blue) or *Enterococcus_D gallinarum* (orange). D) vContact2 network that shows 609 clustered viral metagenome assembled genomes (vMAG) populations present in CBAJ-DB. The network colors represent the vMAG study origin. Pie chart shows proportion of CBAJ-DB vMAGs that clustered to other studies that are cosmopolitan genera (brown), singletons (gray), novel genera (clusters of >1 vMAGs from this study only, green), or known taxonomy (yellow).

We first sought to verify if this microbial dereplicated MAG set represented the key members identified in our amplicon sequenced CBA/J communities from both inflamed (n=14) and uninfected (n=16) individuals (Fig. 1B). The relative abundance of dMQHQ MAGs closely mirrored the full community 16S rRNA amplicon at the class level from both uninfected Day 11 (rho=0.68) and inflamed Day 11 (rho=0.86) mice, indicating the dMQHQ database is representative of CBA/J untreated and inflamed communities (Fig. 3A, Fig. 1C). More specifically, a linear discriminate analysis of MAG relative abundance indicated similar dynamics between our genome and amplicon data sets. For example, *Salmonella* and *Enterococcus_D* were the most significant genomes in determining infected communities, while genomes from *Alistipes, Duncaniella and Lachnospiraceae* COE1 were most significant in determining uninfected communities (Fig. 3B). Additional to these genera, relative abundance of other key taxa is consistent with amplicon sequencing, including *Akkermansia*, and *Muribaculaceae* prominence in uninfected mice and persistence in infected mice. *Lactobacillus* genome and ASV relative abundance also similarly increased during infection (Fig. 3C).

**Fig. 3.**
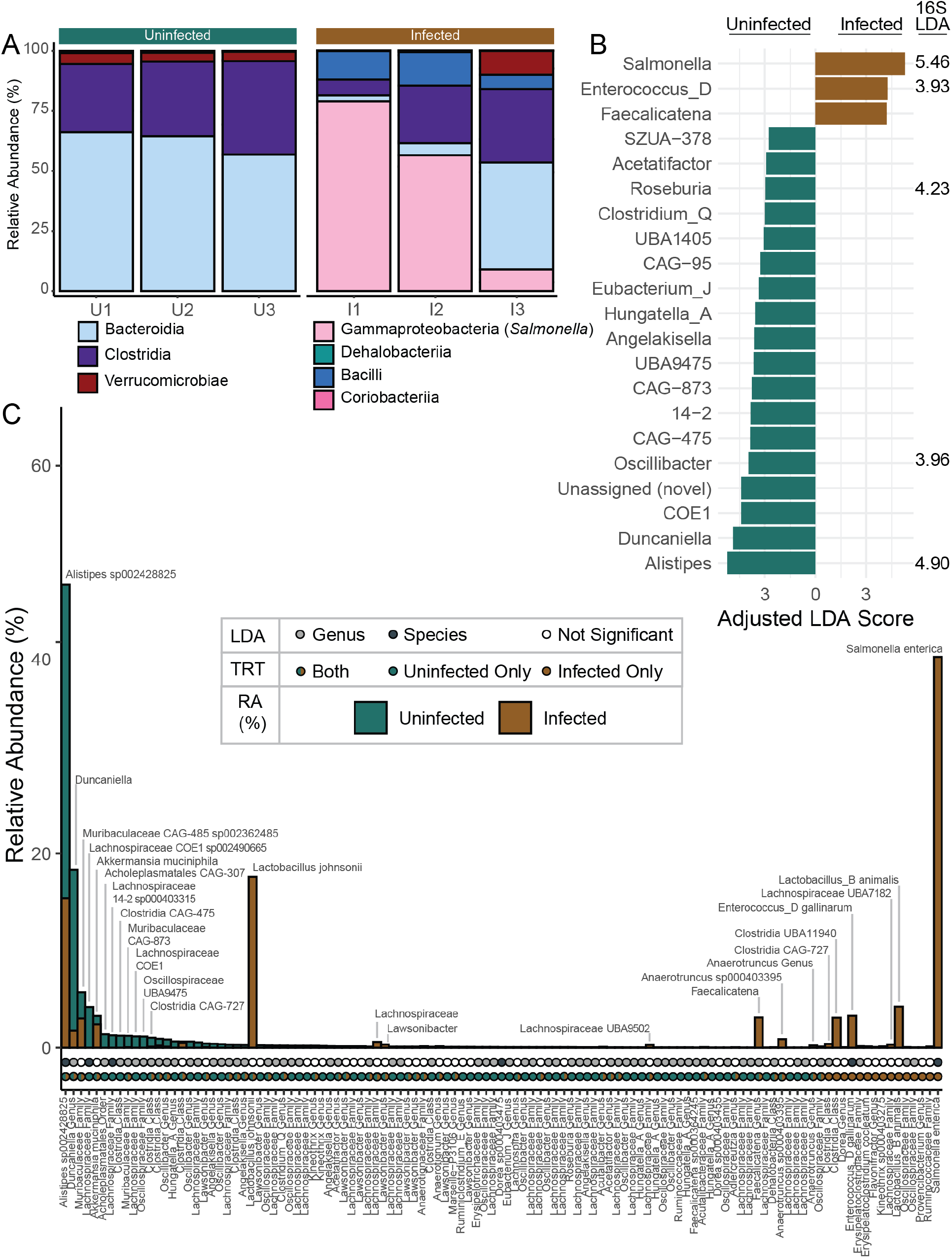
The CBA/J database (CBAJ-DB) genomes are representative key members in amplicon microbiome data. A) Class distribution of dereplicated MQHQ (dMQHQ) metagenome assembled genome (MAG) relative abundance across individual metagenomes (U# = uninfected, I# = infected) labeled by treatment. B) Linear discriminant analysis Effect Size (LEfSe) scores of most important genera in either treatment in metagenomes, including genera LEfSe linear discriminant analysis (LDA) scores for amplicon sequencing variants (ASVs) with matching taxonomy. C) Rank abundance of dMQHQ MAGs (n=113) showing mean relative abundance (RA) in uninfected (green) and infected (brown) treatments (TRT). Circles below bars highlight LDA significant species (black) and genera (gray) (top row) and treatment origin of each MAG (bottom row). MAGs are labeled with the most resolved GTDB-Tk taxonomy.

To link these reconstructed genomes more precisely to the amplicon data, we identified 96 MAGs that contained a partial to full 16S rRNA gene sequence. A pairwise comparison of MAG-derived 16S rRNA sequences and the V4 region sequences from our ASVs identified 33 unique genomes containing sequences matching ASVs in our 16S rRNA dataset. Many MAGs with 16S rRNA matches were among the most enriched taxa including *Lactobacillus johnsonii, Alistipes* sp002428825, and *Clostridia* in order 4C28d-15 (Fig. S3, Data S1). Together these findings indicate significant membership congruence in our MAG database and our amplicon data, demonstrating that inferences made with the CBAJ-DB have relevance to the more broadly sampled amplicon sequenced gut communities from inflamed and uninfected mice.

### This CBA/J microbial genomic resource includes mouse and human relevant lineages

To determine the CBAJ-DB relevance to microbial genomes recovered from other murine models, we compared strain level identity (>99% average nucleotide identity) of our sampled MAGs to similar quality MAGs from two prevalent mouse gut genome catalogs: (i) Integrated Mouse Gut Metagenomic Catalog (iMGMC) and (ii) The Mouse Gastrointestinal Bacteria Catalogue (MGBC). Notably, many of these CBA/J derived genomes represented unique strains from the classes *Bacilli* (n=3), *Bacteroidia* (n=2), *Clostridia* (n=24), *Coriobacteriia* (n=1), and *Dehalobacteriia* (n=1) not represented in iMGMC, and MAGs from *Bacilli* (n=1), *Dehalobacteriia* (n=1), and *Clostridia* (n=30) not represented in MGBC (Fig. 4A). Additionally, of the strains that were sampled in our dataset and prior curated catalogs, 33 (30 *Clostridia*, 3 *Bacilli*) received a higher quality score indicating the value of these recovered MAGs to advance knowledge of cultivated and uncultivated genomes in murine models more broadly.

We also examined CBAJ-DB MAGs against genomes derived from human hosts. To analyze shared genera and species, our dMQHQ database was dereplicated with isolate genomes from the Human Microbiome Project (HMP) (n= 813) and MQHQ MAGs (n=2,560) from a human cohort (PRJNA725020) (Data S5) [31]. *Akkermansia muciniphila* (CBAJDB_482) and *Enterococcus_D gallinarum* (CBAJDB_497), two defining members of the commensal and inflamed gut respectively, clustered with species previously recovered from human hosts. Recovery of *Enterococcus_D gallinarum* from the uninfected CBA/J gut demonstrates the applicability of perturbation techniques to uncover conditionally rare members. As has been reported by others, there was more similarity at higher taxonomic levels (e.g. genus) between our murine and human gut microbial members [32,33], with 27 MAGs from *Bacilli, Bacteroidia, Clostridia, Corriobacteriia, Gammaproteobacteria, and Verrucomicrobiae* sharing similarity (Fig. 4A).

We were particularly interested if the microbial members recovered from our pathogen-inflamed CBA/J had relevance to inflammation in humans. To test this, sequencing reads from the Lloyd-Price et al cohort [34] containing 972 inflamed and 365 healthy gut metagenomes were stringently mapped to the CBAJ-DB MQHQ MAGs (Data S6)[34]. Consistent with their distribution across our treatments, sequencing reads from healthy and inflamed humans mapped to 11 of our *Akkermansia muciniphila* MAGs, while 3 *Enterococcus_D gallinarum* MAGs derived only in our inflamed treatments recruited sequences from inflamed individuals (Fig 4B,C). While it can be challenging to extend specific organismal findings from murine to human conditions [22,32,33], inferences from critical lineages (e.g. *A. muciniphila* or *E. gallinarum)* in our database may have more direct relevance for human-relevant applications.

### *Salmonella* infection and inflammation restructures the metabolic potential of the murine gut microbiome

Given this is one of the first genome-resolved analyses of a pathogen-impacted microbiota, and the first for *Salmonella*, it offered a new opportunity to assess functional potential remodeling during infection. Prior reports indicated that pathogen induced inflammation created oxidative conditions that generated terminal electron acceptors like oxygen, tetrathionate, nitrate, and sulfate [9,19]. As such, we wanted to evaluate the respiratory capacity of inflamed communities and compare it to uninflamed communities. In infected communities, *Salmonella* has the highest mean genome relative abundance, and encoded gene sets for respiring oxygen (both high and low affinity oxidases), fumarate, tetrathionate, and trimethylamine N-oxide (TMAO) (Fig. 5A). Outside *Salmonella*, no other organisms had the capability for respiring with low affinity oxidases, but we infer *Enterococcus and Lactobacillus* have the capability to reduce low levels of oxygen for detoxification (due to the absence of complex I in electron transport chain) while *Akkermansia municiphila* and *Muribaculaceae* likely respire low levels of oxygen using high affinity oxygenase. Similarly, we observed genes for detoxifying reactive oxidative damage (SOD, catalase, thioredoxin reductase) were more enriched in the inflamed community than the uninfected community. Together these findings demonstrated organisms co-existing with *Salmonella* in the inflamed gut encode the metabolic abilities to withstand or leverage the oxidative redox conditions caused by inflammation (Fig. 5C). Markedly, there were members in the uninflamed gut with respiratory metabolic potential that were not maintained in the inflamed gut (*Duncaniella sp, Hungatella_A sp*), demonstrating there are other selective forces besides the ability to respire that dictate persistence in response to pathogen colonization (Fig 5C).

**Fig. 5.**
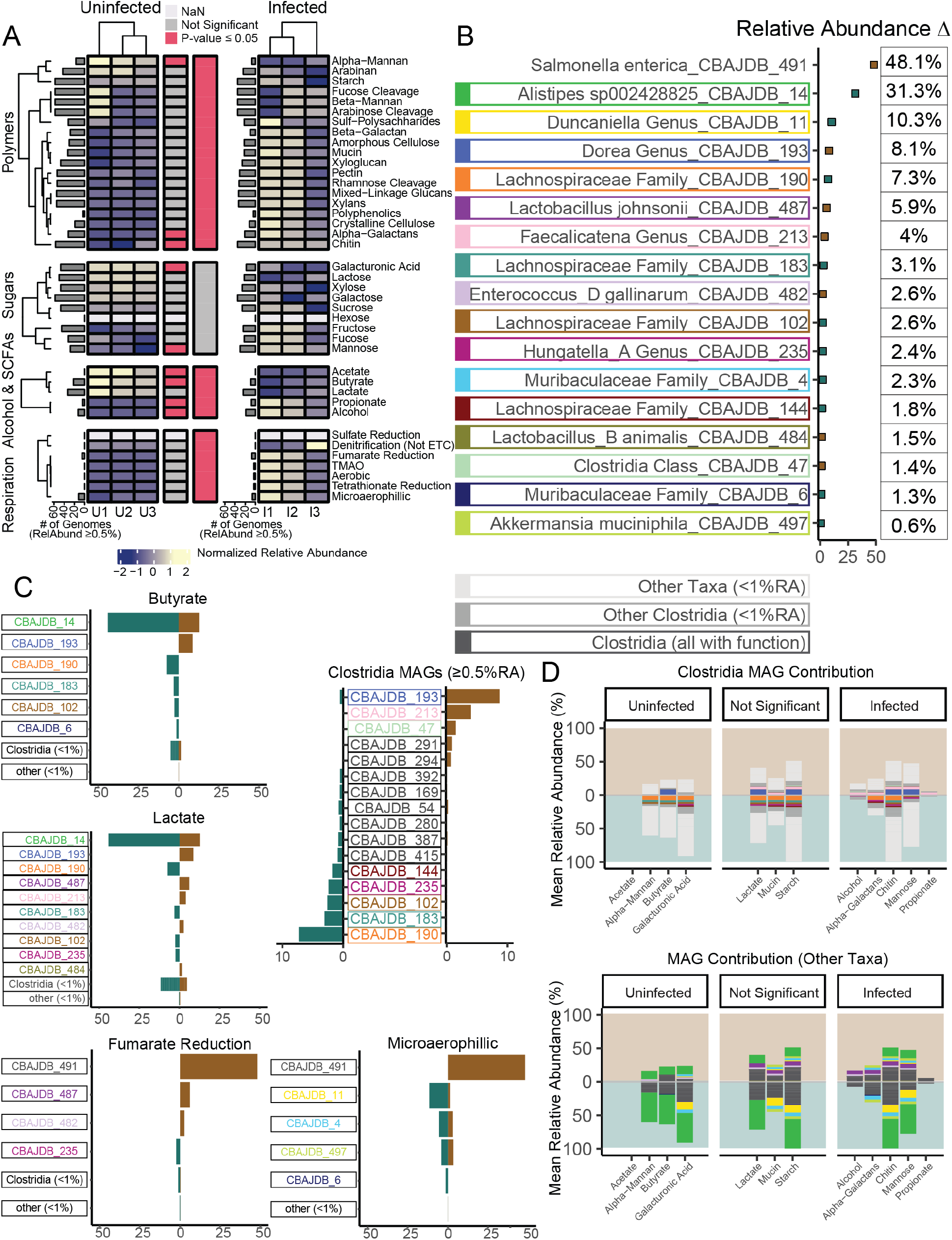
The CBA/J database (CBAJ-DB) highlights differential metabolisms encoded in uninfected and inflamed mice. A) Mean relative abundance of metagenome assembled genomes (MAGs) with each function are shown by a heatmap. Functions that are significantly different between treatments as determined by analysis of variance (ANOVA) (p ≤ 0.05) are indicated by horizontal bars between heatmaps with red highlighting significance at the function (first bar) or functional group level (second bar). Gray bars on each heatmap indicate the number dereplicated medium and high quality (dMQHQ) MAGs that comprise at least 0.5% of the community with a given function. B) Percent change in relative abundance (RA) between treatments the 17 most divergent MAGs. Points are colored by the treatment (uninfected, green; infected, brown) with higher RA. Individual MAGs are uniquely colored by surrounding boxes, acting as color legend for subsequent figure sections. C) Individual MAG contribution to specific functions is shown, with bar magnitude denoting mean RA in either treatment. D) *Clostridia* MAGs with mean RA greater than 0.5% in both treatments are shown. E) *Clostridia* contribution to significant functions (top) and contribution of other taxa (bottom). Plot fields are colored by treatment and bar magnitude indicates mean RA within a treatment.

Prior reports by our team and others demonstrated that butyrate, a key gut short chain fatty acid (SCFA), decreased by 15-fold in the *Salmonella* inflamed gut, most likely due to inflammation induced redox changes with detrimental impacts on members of the class *Clostridia* [13,27]. Here we sought to better understand the relationship between taxonomy and SCFA production potential. In uninflamed communities the most prevalent butyrate producing bacteria were members of the *Alistipes* and *Lachonospiraceae*, members of classes *Bacteroidia* and *Clostridia* respectively. Interestingly, while the most dominant *Clostridia* did decrease in relative abundance with inflammation, replacement *Clostridia* members (*Lachnospiracea, Dorea, Faecalicatena*) were enriched which encoded overlapping butyrate production potential. For example, a MAG belonging to the genus *Dorea* within the *Clostridia* was enriched 16-fold and likely most contributed to butyrate production stability, while the dominant *Alistipes* MAG (a member of the *Bacteroidia*) reduced in abundance by a third was not replaced by taxonomically similar members. Together, these data suggest the notion that decreased butyrate concentrations observed in the CBA/J mouse model during *Salmonella* infection [27] may be attributed to *Bacteroidia* reduction and less so to *Clostridia*, a hypothesis needing further validation using gene expression to track butyrate production and consumption activities in the inflamed gut.

*Salmonella*-induced inflammation alters carbon usage patterns with more favorable redox conditions enabling the use of less energetically favorable substrates like 1,2-propanediol and ethanolamine [35,36]. While *Salmonella* encodes this metabolic capacity, we were interested if any of the other persisting microorganisms could compete for use of these substrates. *Enterococcus_D* and multiple *Oscillospirales* genomes contain genes from the *eut* gene cluster for ethanolamine utilization and *pdu* genes for 1,2-propanediol utilization. These genera increase in relative abundance with inflammation, particularly *Enterococcus_D*, which is one of the next most abundant members after *Salmonella* (expanding to 2.6% of the inflamed community). Additionally, we showed that the polymer utilization profile was also impacted with inflammation, as infected communities can utilize more alpha-galactan and chitin (Fig. 5). In a similar fashion, the community utilization potential of sugars fructose, fucose, and mannose increased with inflammation. Collectively, these data can inform probiotic approaches for controlling *Salmonella* abundance through competitive exclusion targeting select substrate use patterns using inflammation resistant strains.

Next, we quantified genes commonly reported in humans to impact inflammation and examined if they were depleted in this inflamed mouse model. Consistent with literature reporting healthy individuals have a greater potential for tryptophan degradation [37–39], we observed the potential for tryptophanase mediated conversion of tryptophan to indole by members of *Bacteroidia, Clostridia*, and *Verrucomicrobiae* in both inflamed and uninfected mice. However, the proportion of *Bacteroidia* with this gene was much lower in inflamed guts (Fig 6B). Tryptophan Indole/AhR pathway representation in infected mice is concurrent with lower proportions of *Verrucomicrobiae* and *Bacteroidia* spp. (Fig. 6). Also, like human microbiomes, we observed microbial genes responsible for cleaving taurine or glycine from primary bile acids and metabolizing secondary bile acid products (*bsh, baiN, baiA*, and *hdhA*) were significantly lower in relative abundance in mice infected with *Salmonella* (Fig. 6). These data provide promising insights that the functional gene profiles for modulating inflammation may be conserved with humans, supporting CBA/J as a relevant inflammation model.

**Fig. 6.**
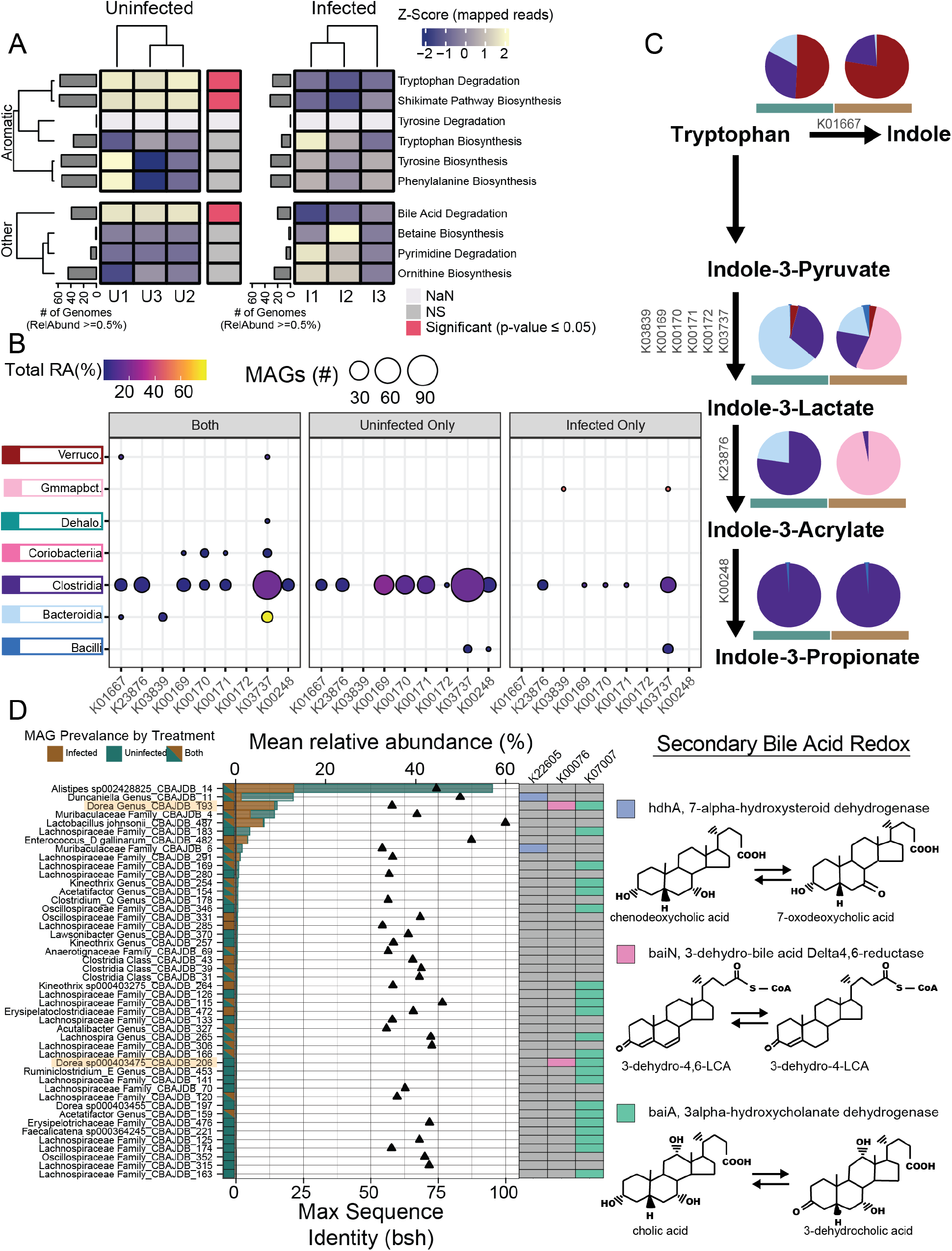
Tryptophan and bile acid metabolism in inflamed and uninfected gut microbiomes. A) Mean relative abundance summed for each function (rows). Functions that are significantly different between treatments as determined by analysis of variance (ANOVA) (p ≤ 0.05) are indicated by horizontal bar between heatmaps with red highlighting significance at the function level. Gray bars indicate the number medium and high quality (MQHQ) metagenome assembled genomes (MAGs) that comprise at least 0.5% of the community with a given function. B) Relative abundance (point color) and number of MAGs (size) in each class with each gene for tryptophan degradation separated by MAG presence in each treatment, where both indicates MAGs that recruited strictly mapped reads from both treatments. C) Tryptophan degradation to indole and indole derivatives pathway with pie charts colored by proportion of MAGs in each class (coloring from B) for each treatment. D) Relative abundance (bars) of MAGs in each treatment encoding bile salt hydrolase (bsh, points show sequence similarity to bsh. or *hdhA* (K22605), *baiN* (K00076), or *baiA* (K07007) involved in secondary bile acid metabolism. *Dorea* are highlighted as MAGs with more than one gene for metabolizing secondary bile acid products.

### Viral AMGs contribute to the bacterial community functional potential in CBA/J mice via *Firmicutes*

In the creation of the first murine gut viral database, we sought to compare viral genomic content cataloged here to other mammalian gut systems. Of the 609 dereplicated vMAGs that were recovered from both treatments, less than 1% had taxonomic assignments (Data S4). These three vMAGs were assigned to the Caudovirales in the families Siphoviridae (n=2), and Myoviridae (n=1). To perform biogeographic analyses, we collated phage genomes previously reported from mammalian guts [24,34,40–42] and clustered these with our mouse recovered vMAGs. We found that 322 of the CBA/J derived vMAGs (53%) had similar representatives in other phage gut metagenome studies, meaning over half of our vMAGs clustered with viruses from at least one additional study (Fig. 4D). This suggests a potentially cosmopolitan phage seedbank that may be conserved across a wide variety of animals, geographies and, in the case of humans, ethnicities and health statuses. Ultimately, viral content in the CBAJ-DB can have relevance to other mouse models and human guts.

To explore if viral communities could potentially influence the structure and function of the CBAJ-DB uninflamed and inflamed microbial communities, we verified that microbial and viral genome-based ordinations were coordinated (Fig. S4). With informatics we conservatively determined that of the 609 vMAGs, 11.5% were putatively linked to 43 MAGs that encompassed 27 unique taxonomies (Fig. S4). All putative hosts corresponded to members of the *Firmicutes*, and included members of the *Lachnospiraceae, Ruminococcaceae, Oscillospiraceae, Anaerotignaceae, Borkfalkiaceae* and *Acutalibacteraceae* families. Among the vMAGs that putatively infected hosts, we identified 36 auxiliary metabolic genes (AMGs) with functionalities including regulation of the TCA cycle (citrate synthase), glycolysis (orthophosphate dikinase), phosphate metabolism (PhoH), and oxidative stress response (rubrerythrin). These phage genomes also encoded AMGs for the induction of germination (Peptidase A25), spore formation (M50B), the cleavage of amorphous cellulose (GH2), and low pH resistance (ornithine carbamoyltransferase). Among the putative viral hosts were members within the *Clostridia* class, exhibiting some of the largest MAG relative abundance differences between inflammation states. For example, *Dorea* and *Faecalicatena* enriched in inflamed mice, and *Lachnospiraceae* COE1 enriched in uninfected mice. Together these findings indicate phages may be underappreciated top down (predation) and bottom up (resource) controllers of microbiota functionality in the murine gut.

## Discussion

### Perturbation expanded the microbial and viral genomic cataloging of the CBA/J gut microbiome

Genome resolved catalogs like CBAJ-DB are valuable resources for interrogating metabolic potential of the microbiome, yet these databases are biased by the by environment, organism, or disease state they were generated from; and host associated microbiomes can vary drastically between different species, model organisms, and even within an individual [43–45]. Previous murine gut bacterial databases lacked membership from inflamed individuals and CBA/J mice, and none have curated viral content [21,22,46], underscoring a clear value of this resource to the community. Our findings highlight the power of using biological perturbation, in this case *Salmonella-*induced inflammation, to genomically sample taxa that are conditionally rare and obscured by their low abundance only to be critical contributors to ecosystem functionality under altered states.

While assemblies and binning are prone to missing key lineages or under-sampling diversity, we used paired 16S rRNA analyses to affirm the representation of critical community members in the inflamed and uninfected gut. Our paired amplicon sequencing indicates the CBAJ-DB contains membership similar to previously reported *Salmonella*-inflamed CBA/J communities and proportional representation of similar bacteria to those found in other mouse breeds [13,27,47]. It is our intent to create a resource for other researchers conducting microbiome analyses using CBA/J mice, such that this genome content can be accessed by taxonomic naming or linkages to the 16S rRNA gene. Likewise, this genome library can be used by others for read recruitment of future metagenome and metatranscriptome sequencing, or to substantiate metabolomic insights from the CBA/J microbiome.

The vMAG database also provides interesting context for researchers in other mammalian gut habitats, but especially mouse gut which has been historically under-sampled in this regard. While collections of human gut viruses are available, the mouse virome is understudied [24,48]. This collection of over 4,000 vMAGs contains a significant number of cosmopolitan genera also found in other mammalian guts including humans. The existence of a core mammalian gut virome is an exciting proposition that alludes to an intricacy in gut community function and begs further exploration. The gut microbiome is a complex system involving the interplay of host, microbes, and abiotic elements [49–51]. Beyond functional characterization of gut communities during *Salmonella* infection, the CBAJ-DB offers a bacterial and viral resource for holistic microbiome study in a common mouse model.

### A genome resolved inventory of functional potential changes in the pathogen inflamed gut

Previous studies of gut microbiomes highlight the role microbial metabolites play in gastrointestinal inflammation as signaling effectors in host immune regulation [52,53]. The CBAJ-DB showcases the juxtapose of pro-inflammatory and anti-inflammatory microbial membership and gene content in CBA/J mice and during *Salmonella* infection. *Salmonella*-induced inflammation shifted the functional potential of infected communities favoring respiring organisms, marked by a reduction of butyrate and acetate producers and an increase in bacteria with anaerobic respiration capability.

Convention indicates butyrate agonism of PPAR-γ and SCFA engagement with G-protein coupled receptors respectively help to maintain luminal anaerobiosis and promote colonic regulatory T cell development [6,54,55]. We showed specific bacteria reduction coincides with an inflamed state and diminished SCFA production potential in the gut.

Our findings indicate *Alistipes* reduction following inflammation as a possible cause of butyrate production potential loss, contrasting with current dogma linking butyrate production in the gut chiefly to *Clostridia* abundance [11,56–58]. Furthermore, *Salmonella* infection enriched certain *Clostridia* including *Dorea, Faecalicatena*, and a novel bacteria described only at the class level, highlighting the functional redundancy that may be provided within this class. Interestingly, the mouse with the lowest *Salmonella* relative abundance had the greatest diversity of *Clostridia*. It is interesting to speculate that this lineage and the diversity within it, may be important for microbiota recolonization and return to homeostasis following gastric infection, a notion supported by previous research [59].

*Salmonella-*induced inflammation also caused a reduction in bacteria with the capability to mediate anti-inflammatory microbial metabolites. Bacteria with genes for secondary bile acid production and tryptophan catabolism were decreased in inflamed metagenomes, a response previously shown to increase host susceptibility to infection [60]. Specifically, reduced bile acid can limit ligands like pregnane X (PXR) and farnesoid X (FXR) which are important regulators of the host anti-inflammatory response, thus reduced production of these genes could have further feedback on inflammation [64,65]. Similarly, indole and indole derivatives like indole acrylic acid, indole-3-probionate, and indole-3-lactate are AhR and PXR agonists and thus anti-inflammatory [37,61]. Here we demonstrate how the reduction of bacteria capable of producing these important host pathway modulators can further promote inflammation and *Salmonella* expansion, evidenced by high lipocalin-2 levels concomitant with high *Salmonella* relative abundance and lower functional potential for bile acid deconjugation, secondary bile acid production, and tryptophan catabolism in inflamed mice. Future research directly measuring transcription and metabolite concentrations can be used in concert with the CBAJ-DB to determine the anti-inflammatory impact of individual bacteria on the microbiome.

### Commensal bacteria that can withstand inflammation may represent future biological therapeutic opportunities

A recent rise of antibiotic resistant *Salmonella* strains globally underpins the need for alternative treatments and prevention measures against foodborne pathogens. One avenue may be the use of probiotic bacteria to reduce the intensity or duration of infection [37,62,63]. *Alistipes* sp002428825 and *Akkermansia muciniphila* clustered closely with genomes from human microbial communities and we explore here their potential as anti-inflammatory probiotics.

*Akkermansia muciniphila* is a well-known commensal gut bacterium in mammals that lives in the lumen mucosal layer and contributes to epithelial gut barrier integrity [64,65]. Nevertheless, one study showed *Akkermansia muciniphila* exacerbated inflammation and increased *Salmonella typhimurium* relative abundance [66]. These findings are inconsistent with our data however, where *Akkermansia muciniphila* is relatively abundant and consistently present in uninfected and *Salmonella*-infected mice. Genome analysis reveals the capacity of *Akkermansia muciniphila* to produce indole, potentially an important anti-inflammatory mechanism and *Salmonella* deterrent. Other studies have shown the effectiveness of indole from *E. coli* increasing tight junctions of the gut epithelial and decreasing *Salmonella* pathogenicity, though more work is needed to confirm similar action from *Akkermansia* in buffering gut inflammation [67].

*Alistipes* sp002428825 was also consistently detected in both treatments. Analysis of this genome suggests it can respire oxygen and directly compete with *Salmonella* for arabinan, arabinose, and pectin, whilst maintaining critical gut homeostasis functionalities like butyrate production. *Alistipes* spp. are often associated with healthy human microbiomes [68,69] and have even been shown to facilitate microbiota recolonization following perturbation [70]. We have shown *Alistipes* has saccharolytic genes to metabolize rhamnose and fucose and it is possible that Gram-positive bacteria eliminated by inflammation and fucosylated proteins from host epithelial erosion could support *Alistipes* during *Salmonella* infection as sources of these sugars [71,72]. Given this genus, like *Akkermansia muciniphila*, remained abundant despite host inflammation, and both bacteria contain genes for indole derivative production, it also may be directly antagonistic to *Salmonella* through anti-inflammatory effector potential.

It also bears mentioning the presence of lactic acid bacteria *E. gallinarum* and *L. johnsonii* in the CBAJ-DB in relatively high abundance during *Salmonella* infection. In fact, these members were previously illuminated in prior studies from mice as well as clinical patients, demonstrating co-occurrence may exist beyond this single study or mouse model [73]. Beyond resisting or maybe even responding to conditions created by *Salmonella* infection, we provide genomic evidence supporting *L. johnsonii* nutrient competition for arabinose, mixed-linkage glucans, and amorphous cellulose, while *E. gallinarum* may compete for chitin, pectin, and arabinan. Additionally, closely related species to these have been shown to produce anti-*Salmonella* agents like bacteriocin and organic acids putatively decreasing *Salmonella* abundance over time [74–76]. Given these species have many members already approved as probiotics and our data indicate CBA/J mice harbor at least one species of *Enterococcus* common to human guts and *Lactobacillus* spp. resistant to host inflammation, a probiotic lactic acid bacteria strain resistant to *Salmonella* may reside in the CBAJ-DB [77]. This genome resolved research identifies future targets with promising potential for exploration as probiotics robust to *Salmonella* infection. Yet, we recognize that commensal bacteria like these can be pathobionts provided the right setting [43,66,78,79], such that future research in this mouse model would first include challenge experiments with isolates or even targeted consortia that provide multiple avenues of overlapping pathogen colonization resistance.

### Conclusion

The CBAJ-DB uniquely captures gut community variation in CBA/J mice. MAG reconstruction from metagenomic sequencing enabled us to profile the functional potential of the murine gut microbiome during acute *Salmonella* inflammation, contrasting community membership and gene content with uninfected mice. Persisting taxa in the inflamed gut encoded the capability to withstand or utilize changing redox conditions, while bacteria producing SCFA and producing host anti-inflammatory effectors decreased in mice with high *Salmonella* burden. Further, our phage analyses leave open the possibility that phage infection could alter *Firmicutes* energy regulation and spore formation.

Together, these findings validate physiological investigations performed with reduced complexity synthetic or modified gut microbiota. We also provide new perspectives that advance the understanding of *Salmonella* effect on an intact microbiome and provide model-specificity to CBA/J gut consortia. Our efforts show novel bacteria unique to the CBA/J mouse model and enriched by *Salmonella* infection. An exploration of potential probiotic targets in the CBAJ-DB revealed multiple lactic acid bacteria capable of withstanding the host immune response to *Salmonella* and that may be indifferent to *Salmonella* competition. Additionally, genomes were recovered of *A. muciniphila* and *E. gallinarum* with species similarity to bacteria in human guts. The CBAJ-DB is the first culture-independent murine genome catalog to include sampling from CBA/J mice and inflamed individuals, providing a resource with application to multi-omic microbiome investigation, gut inflammation research, and studies involving the CBA/J mouse model broadly.

## Methods

### Strains and media

*S. enterica* serovar Typhimurium strain 14028 (*S. typhimurium* 14028) was cultured overnight in Luria-Bertani (LB) broth at 37 °C with constant agitation. Overnight culture was washed and resuspended in water. *S. typhimurium* 14028 ASV was determined manually from an identical sequence match to the 16S region of NCBI Reference Sequence NZ_CP034230.1 mapped with Geneious Prime® 2020.1.2.

### Animals and experimental design

Female CBA/J mice were purchased from The Jackson Laboratory (Bar Harbor, ME) and housed 5 per cage in conventional enclosures maintained in a temperature controlled 12 hour light/dark cycle. Irradiated mouse chow (Teklad, 7912) was made available *ad libitum* to mice in the control group (n=16) and to infected mice (n= 14) housed separately. Individuals in this study were chosen based on fecal sample availability at Day 11 and only infected mice with ≥25% *S. typhimurium* 14028 at the sampling time were used. Mice in the infected group received 10^9^ CFU *S. typhimurium* 14028 oral gavage on Day 0 with no subsequent treatment, and control group mice were left without treatment. Animal experiment protocol was approved by The Ohio State University Institutional Animal Care and Use Committee (IACUC; OSU 2009A0035).

### Sample collection

Fecal pellets were collected from each mouse one day prior to treatment and 10 and 11 days after treatment initiation on autoclaved aluminum foil. Pellets were immediately placed in labeled microcentrifuge tubes and flash frozen with EtOH/dry-ice prior to storage at −80 °C until further processing.

### Lipocalin-2 quantification

Vortex homogenization of fecal sample in PBS containing 0.1% Tween 20 (100 mg/ml) for 20 min was performed prior to centrifugation of the resultant suspension at 12,000 rpm for 10 min at 4 °C. The resulting supernatant was used to measure levels of inflammation marker Lipocalin-2 using the Duoset murine Lcn-2 ELISA kit (R&D Systems, Minneapolis, MN).

### DNA extraction and sequencing

Total nucleic acids were extracted using the *Quick-*DNA Fecal/Soil Microbe Microprep Kit (Zymo Research) and stored at −20 °C until amplicon or metagenomic sequencing. DNA was submitted for amplicon sequencing at Argonne National Lab at the Next Generation Sequencing facility using Illumina MiSeq with 2 × 251 bp paired end reads following established HMP protocols [80]. Briefly, universal primers 515F and 806R were used for PCR amplification of the V4 hypervariable region of 16S rRNA gene using 30 cycles. The 515F primer contained a unique sequence tag to barcode each sample. Both primers contained sequencer adapter regions. DNA for metagenomes was submitted to the Genomics Shared Resource facility at Ohio State University and was prepared for sequencing with a Nextera XT library system followed by solid-phase reversible immobilization size selection. Libraries were quantified and then sequenced using an Illumina HiSeq platform.

### 16S rRNA preprocessing

Amplicon sequencing fastq data were processed in a QIIME2 2019.10.0 environment [81]. Reads were demultiplexed and then denoised with DADA2 [82]. For all sequencing runs (n=4), forward reads were truncated at 246 bps and reverse reads were truncated at 167 bps. Feature tables from each sequencing run were combined and ASVs were assigned taxonomy with the silva-132-99-515-806-nb-classifier [83]. Before further analysis the ASV table was filtered with R version 4.0.2 to (i) remove samples with no ASVs, (ii) to remove samples with a combined ASV count <1000 across all samples, (iii) to remove ASVs with 0 abundance in every sample, and (iv) to remove ASVs designated as mitochondria and chloroplast. The resulting filtered feature table contained 23,022 ASVs. Raw reads are deposited on NCBI (PRJNA348350) and the final ASV table is published in the supplementary materials (Data S1).

### Genome reconstruction from metagenomes

Quality scores of raw metagenomic reads were evaluated using FastQC (v0.11.9, [84]). Reads were trimmed, adapters removed, and mouse reads removed using BBDuk from BBTools (v38.89, https://jgi.doe.gov/data-and-tools/bbtools). Trimmed reads from each individual sample were assembled with Megahit (v1.1.1, [85]) and with IDBA_UD (v1.1.3, [86]). Each assembler was also used to perform co-assembly of all the samples (n=6) at once.

Subsequently, each single-sample and co-sample assembly was binned separately. To obtain bins, reads were mapped to assembly contig or scaffold set filtered to ≥2500 bps using BBMap (https://jgi.doe.gov/data-and-tools/bbtools), sorted using SAMtools (v1.9, [87]). and then binned with Metabat2 (v2.12.1, [88]). The resulting MAGs were checked for quality and contamination with CheckM (v1.1.2, [89]). Using the resulting medium quality (completeness ≥ 50%, contamination < 10%) and high-quality (completeness ≥ 90%, contamination < 5%) MAGs from the initial single-sample and co-sample assemblies, trimmed reads were mapped using BBMap in perfect mode. Reads that did not map were assembled individually by-sample and co-assembled by treatment with IDBA_UD (v1.1.3, [86]) to create subtractive assemblies. The resulting subtractive contigs or scaffolds were then subject to previously described filtering (≥2500 bps), processing, binning, and quality check.

To construct the final CBAJ-DB, all MQHQ MAGs from all assemblies were assigned taxonomy with GTDB-Tk (v1.3.0, r95, [90]) and dereplicated with dRep (v2.5.4, 99% ANI, [91]). Conventional assembly and binning of reads rarified with BBTools for MAG recovery sensitivity to read depth was performed with the Snakemake [92] workflow included here (https://github.com/ileleiwi/metaG_pipeline/blob/main/Snakefile_5.88_onesamp).

### 16S rRNA linked to MAGs

Mining of MAGs for 16S rRNA genes was performed with Wrighton Lab software (https://github.com/WrightonLabCSU/join_asvbins) MMseqs2 (v13.45111, [93]) and Barrnap (v0.9, [94]) and SILVA reference database (SILVA_138_SSURef_NR99_tax_silva, [83]) followed by pairwise comparison to the V4 region sequences from our amplicon sequencing.

### Database mapping and comparison to other MAG resources

To calculate relative abundance of individual dMQHQ MAGs, BBmap (https://jgi.doe.gov/data-and-tools/bbtools) was used to randomly map trimmed reads to the dMQHQ with minid = .95. Then CoverM (v0.6.0, https://github.com/wwood/CoverM) was used to estimate read counts per scaffold and per bin. Values are the mean number of aligned reads calculated after removing positions with the most and the least coverage as determined by default values (methods: trimmed_mean, covered_fraction, reads_per_base). Relative abundances of mapped reads were then GeTMM normalized (Data S2) [95] using the edgeR package (v3.36.0, [96]). To place MAGs in either treatment (control, infected) or both treatments, CoverM was used and mapping was only considered if the subject covered fraction exceeded 75% with 95% sequence identity and a minimum of 3 reads per base depth.

To estimate the relative abundance of each vMAG, the metagenomic reads were mapped using Bowtie2 [97]. Reads were mapped using BBMap with minid = 0.95. Afterwards, CoverM was run using the - mean option to consider only those vMAGs that have >75% of their fraction covered. Relative abundances for each vMAG were calculated as their coverage proportion from the sum of the whole coverage of all bins for each set of metagenomic reads prior to GeTMM normalization and are reported in Additional file Data S4.

Isolate MAGs were obtained from the Human Microbiome Project [31] and medium and high-quality MAGs were obtained from https://doi.org/10.5281/zenodo.4977712. Metagenomic reads from the Human cohort and Lloyd-Price cohort [34] were obtained from PRJNA725020 and the SRA Database Commons respectively. Human cohort rarefication to 2Gbps was performed with BBTools reformat. Human metagenomic reads were mapped to MQHQ bins with fastANI (v1.32, [98]). Genome matches with ANI ≥ 94% and alignment fraction (AF) ≥ 33% were deemed of the same species and genome matches with ANI ≥ 73% and AF ≥ 33% of the same Genus [99].

Comparison to mouse database MAGs was performed on MGBC [22] non-redundant high-quality genomes (Data S5, MGBC-hqnr_26640) and iMGMC [21] dereplicated medium and high-quality MAGs (Data S5, iMGMC-mMAGs-dereplicated_genomes). iMGMC MAGs were re-classified with GTDB-Tk version 1.3.0 r95 to align with CBAJ-DB and MGCB taxonomy, and dereplication with each MAG set and CBAJ-DB was performed with dRep (v2.5.4, 99% ANI).

### MAG function analysis

CBAJ-DB MAGs were annotated using DRAM (v1.1.1) [30] and dbCAN (v3.0.2, dbCAN-HMMdb-V10, [100]). CAZy genes called with DRAM were updated with the latest dbCAN database and parsed with HMMer (v3.3, [101]) to include only significant hits (e-value < 10^−18^) with > 35% coverage. Wrighton lab software rule_adjectives (https://github.com/rmFlynn/rule_adjectives) and functions_pa (https://github.com/ileleiwi/functions_pa) were used to parse KEGG ids, Enzyme Commission numbers, and dbCAN ids from gene annotations referencing function rule sheets (Data S7) to determine function presence or absence in each MAG. Function relative abundance significance was determined with ANOVA performed using the R stat package (v4.1.3).

### Viral host-linkage and AMGs

Metagenomic assemblies and subassemblies (n=16) were screened for DNA viral sequences using VirSorter2 (v2.2.2, [102]) using the published protocol in Protocols.io [103]. Briefly, VirSorter2 was run using parameters “–include-groups dsDNAphage, ssDNA”, “--min-length 10000”, and “–min-score 0.5”. The resulting VirSorter2 output was then run through CheckV [104] to ensure quality viral sequences using the “end_to_end” function. The trimmed viral sequences output by CheckV were then once again run through VirSorter2 using options above with additional flags “--seqname-suffix-off” “--viral-gene-enrich-off”, and “--prep-for-dramv”. The final output was then manually curated using the CheckV output as described in Protocols.io. Briefly, 1) viral-like scaffolds that had more than 1 viral gene were kept and deemed viral 2) viral-like scaffolds with no viral genes, host genes equal to zero, or score ≥0.95, or that had 1 host gene with a length of ≥10kb were separated and further inspected. Scaffolds not meeting the above criteria or manually inspected to be non-viral were discarded. After generation of a curated, quality-controlled viral vMAGs, they were clustered at 95% identity across 85% of the shortest contig representing viral populations using ClusterGenomes [103].

To determine taxonomic affiliation, vMAGs were clustered to viruses belonging to viral reference taxonomy databases NCBI Bacterial and Archaeal Viral RefSeq V85 with the International Committee on Taxonomy of Viruses (ICTV) and NCBI Taxonomy using the network-based protein classification software vConTACT2 (v0.9.8, [105]) using default methods [106]. To determine if the viruses present in this study represented relevant communities across mammalian gut ecosystems, we included viruses mined from 5 publicly available datasets in our vConTACT2 analyses from: 1) human guts [24,34,40], moose rumen [41], and bird / other mammalian guts [42]. The viral sequences that were identified from these systems and the genes used for vConTACT2 are deposited on Zenodo with doi ##.####/zenodo.####### with more information of downloaded datasets found in Additional file Data S4.

Viral contigs were annotated with DRAM-v [30]. Auxiliary scores were assigned by DRAM, based on the following ranking system: A gene is given an auxiliary score of 1 if there is at least one hallmark gene on both the left and right flanks, indicating the gene is likely viral. An auxiliary score of 2 is assigned when the gene has a viral hallmark gene on one flank and a viral-like gene on the other flank. An auxiliary score of 3 is assigned to genes that have a viral-like gene on both flanks. All vMAG annotations are reported in Additional file Data S4. To identify likely vMAG hosts, we used two strategies which included 1) Linking viral spacers found in CRISPR systems assembled using CRASS [107] and 2) Oligonucleotide frequencies between virus and hosts using VirHostMatcher and a threshold of d2* measurements of <0.25 [108]. The lowest d2* value for each viral contig <0.2 was used, and only vMAGs for which the top 3 hits had taxonomic consensus at the genera level were considered “good” hits [108]. All virus-host links are reported in Additional file Data S4.

### Spearman Correlation of Metagenomic and Amplicon Communities

Relative abundance of mapped reads to dMQHQ MAGs were averaged within treatment and the total relative abundance of each MAG in *Bacteriodia, Clostridia, Verrucomicrobiae, Gammaproteobacteria, Bacilli*, and *Coriobacteriia* Classes was summed respectively. Relative abundance of MAGs from all other classes were summed together into an additional category. Similarly, 16S amplicon sequence ASV relative abundances from high responder mice and control mice were summed within each treatment and by taxonomy according to the classes previously mentioned, combining all other ASV abundances into an additional category. Spearman correlation was then performed with the R stats package comparing Class abundances between metagenomic communities and ASV communities within the same treatment.

### Statistical analysis

Alpha (Shannon’s diversity) and Beta diversity (Bray-Curtis dissimilarity) were calculated with ASV or MAG relative abundances from the filtered data using the Vegan package 2.5.7 in R [109]. The NMDS Beta diversity visualization was produced using ggplot in R [110]. To determine significant grouping of samples by treatment, we performed an analysis of similarities and mrpp [111], with a stress test determining goodness of fit [111]. Lipocalin-2 (ng/g) and *S. typhimurium* 14028 relative abundance significance between treatments was determined using a Wilcoxon Rank Sum Test, the same test used to compare class relative abundance between different treatments and timepoints. Linear discriminate analysis on MAGs and ASVs was done with LEfSe [112]. MAG and vMAG ordination coordination was determined by Procrustes analysis [109].

## List of abbreviations

CBAJ-DB: The CBA/J database
MAG: Metagenome assembled genome
vMAG: Viral metagenome assembled genome
ASV: Amplicon sequencing variant
NTS: Non-Typhoidal *Salmonella*
MQHQ: Medium and high-quality
dMQHQ: Dereplicated medium and high-quality
AMG: Auxiliary metabolic gene
iMGMC: integrated Mouse Gut Metagenomic Catalog
MGBC: Mouse Gastrointestinal Bacterial Catalogue
PXR: Pregnane X receptor
FXR: Farnesoid X receptor
AhR: aromatic hydrocarbon receptor
SCFA: Short chain fatty acid

## Declarations

### Ethics approval and consent to participate

Mouse experiments in this study were performed in accordance with protocols approved by The Ohio State University Institutional Animal Care and Use Committee (IACUC; OSU 2009A0035-R4).

### Consent for publication

All authors consent to the manuscript submission to *Microbiome*.

### Availability of data and materials

The sequence data supporting the results of this article are available in the National Center of Biotechnology Information (NCBI) under Bioproject number PRJNA348350. All sequencing outputs including the ASV table and fasta files are included on Zenodo ##.####/zenodo.#######.

## Authors’ contributions

IL, MS, JRR, and MAB analyzed and interpreted sequencing data. ASD, IL, MS handled mice and sample collection and ASD performed lipocalin-2 assays. RAD was responsible for DNA extractions and quality control. KK, LK, and LMS provided conceptual framework and editorial support. RMF and IL wrote function parsing scripts and MAB and KCW provided function rules. ASD and BA handled *Salmonella* aspects of the project.

***Acknowledgements***

The authors would like to thank Tyson Claffey and Richard Wolfe for Colorado State University server management; Sandy Shew for management of computing resources retained from The Ohio State University Unity cluster; and Dr. Pearlly Yan at the Genomics Shared Resource Core at The Ohio State University Comprehensive Cancer Center for management of metagenomic sequencing

---

**Fig. S1.**
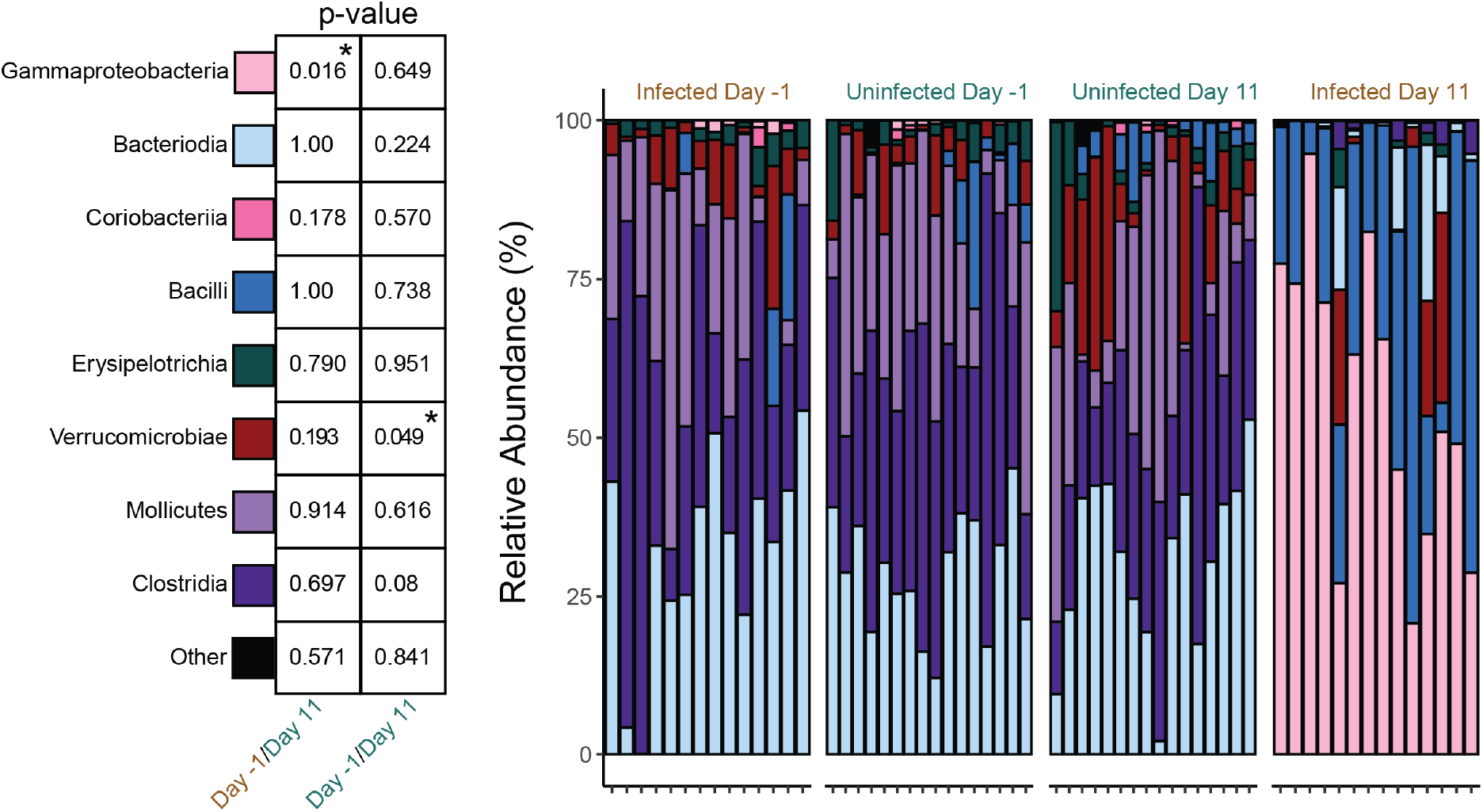
Relative abundance of classes in pre-infection communities are no different than uninfected communities. Wilcoxon Rank Sum significance revealing no difference between Infected Day −1 communities and uninfected communities from either Day −1 or Day 11. Summed abundance within each class was examined comparing different timepoints and treatments (uninfected = green, infected = brown).

**Fig. S2A.**
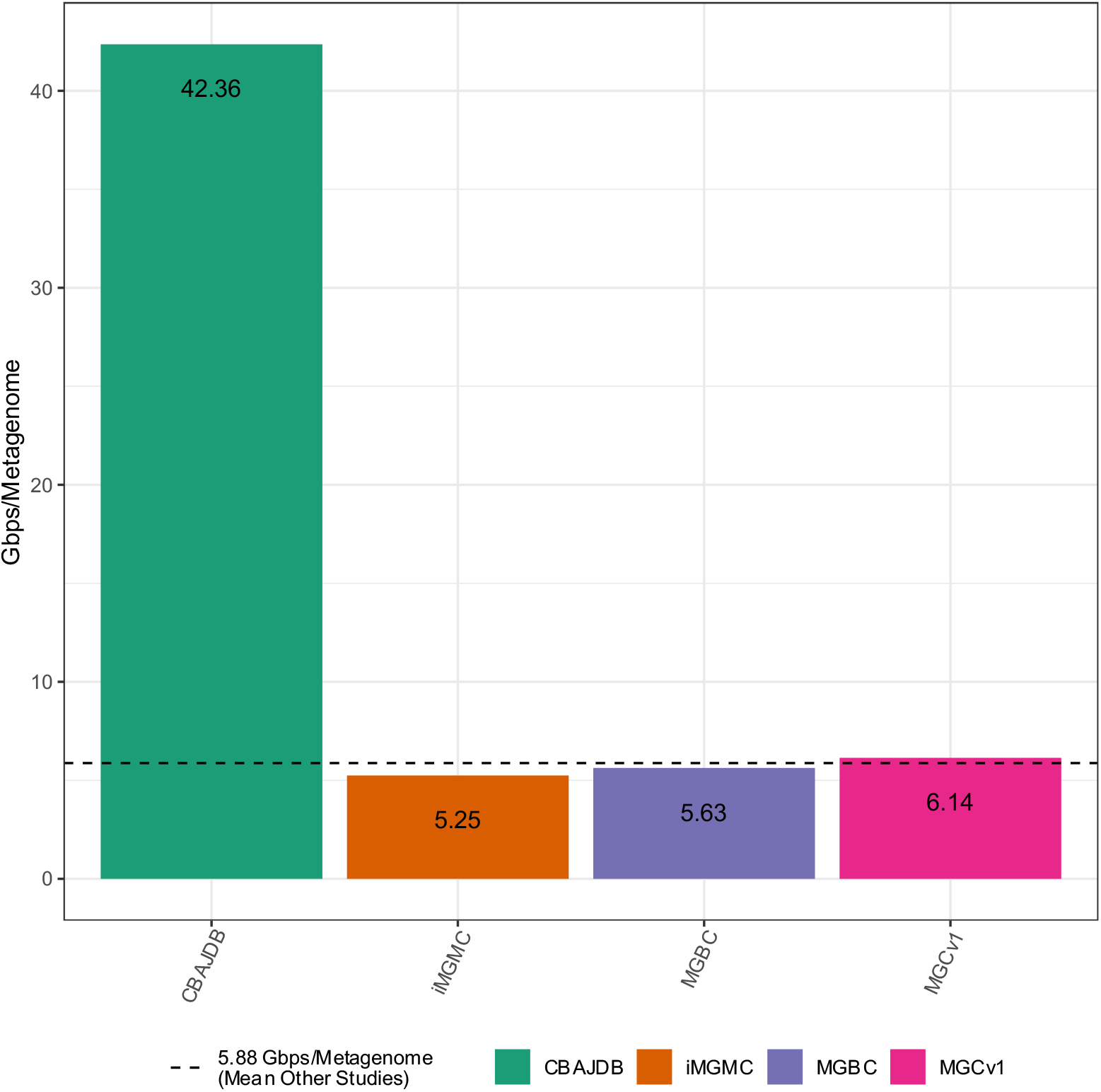
Gigabase pairs (Gbps) per sample of prevalent murine genome databases. integrated Mouse Gut Metagenomic Catalog (iMGMC), The Mouse Gastrointestinal Bacterial Catalogue (MGBC), and Xiao et.al. 2015 (https://doi.org/10.1038/nbt.3353) (MGCv1) compared to the per sample sequencing effort of the CBAJ-DB.

**Fig. S2B.**
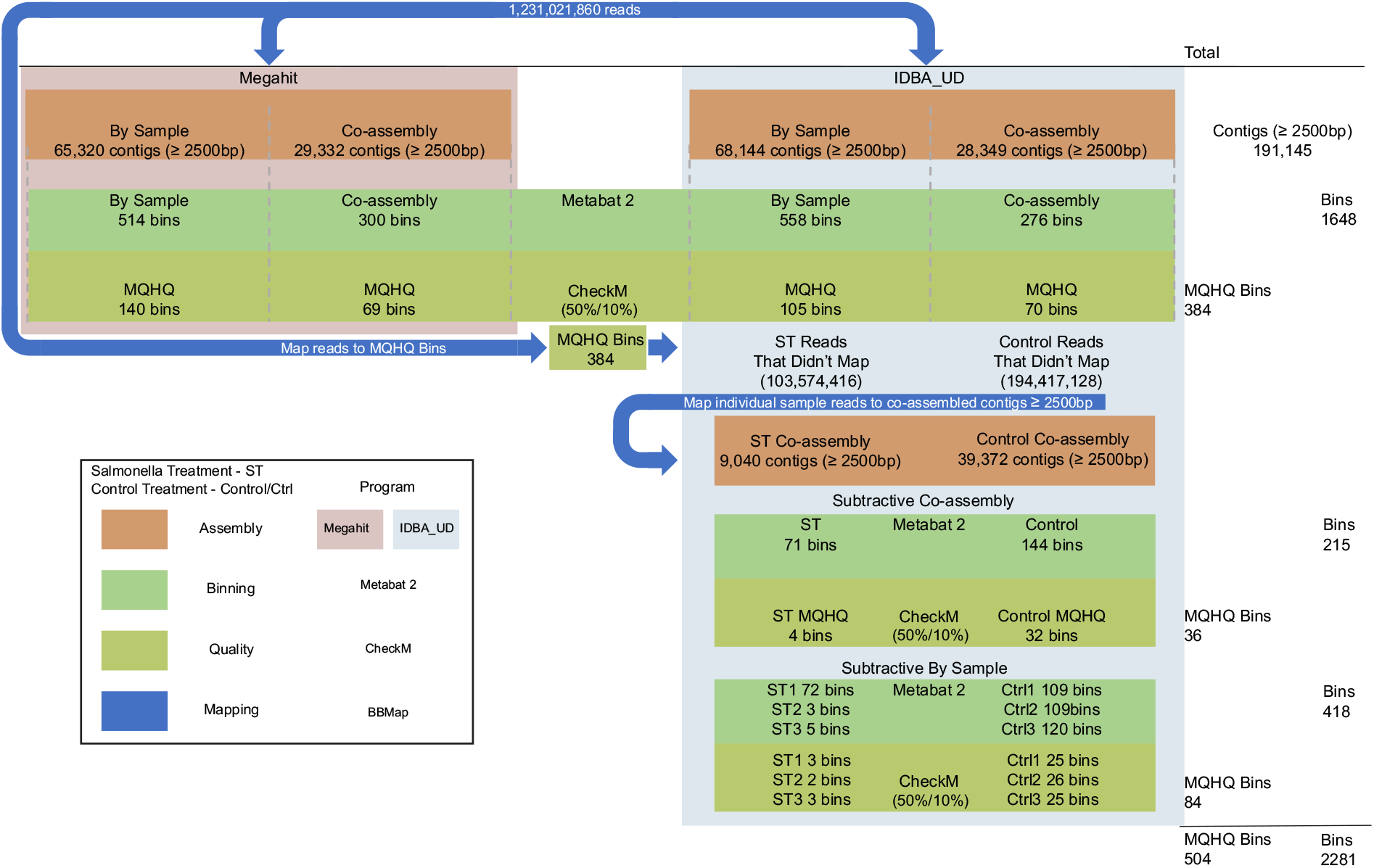
Assembly and binning workflow. Total and medium and high quality (MQHQ) bins derived from Megahit assembler (left) and IDBA_UD assembler (right) shown, blue arrows indicate read mapping for subtractive assembly.

**Fig. S3.**
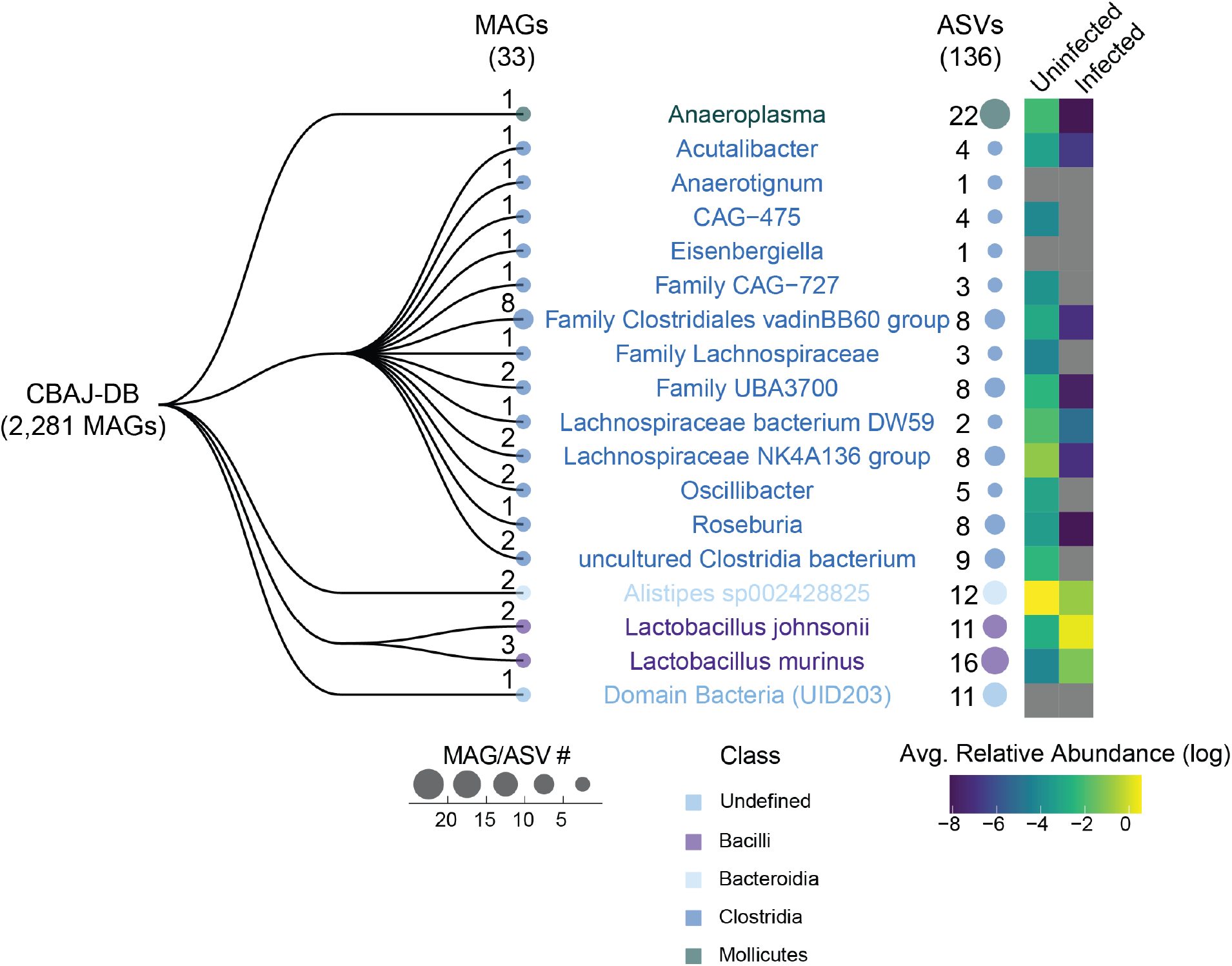
Most resolved taxonomy groups of metagenome assembled genomes (MAGs) containing amplicon sequencing variants (ASVs). MAG groups with matching ASVs from 16S rRNA sequencing as determined by Mmseqs2 and Barrnap (see methods). Colored text indicates the lowest resolved taxonomy. Mean ASV relative abundance within each taxonomy group in each treatment is represented in Log relative abundance (right).

**Fig. S4.**
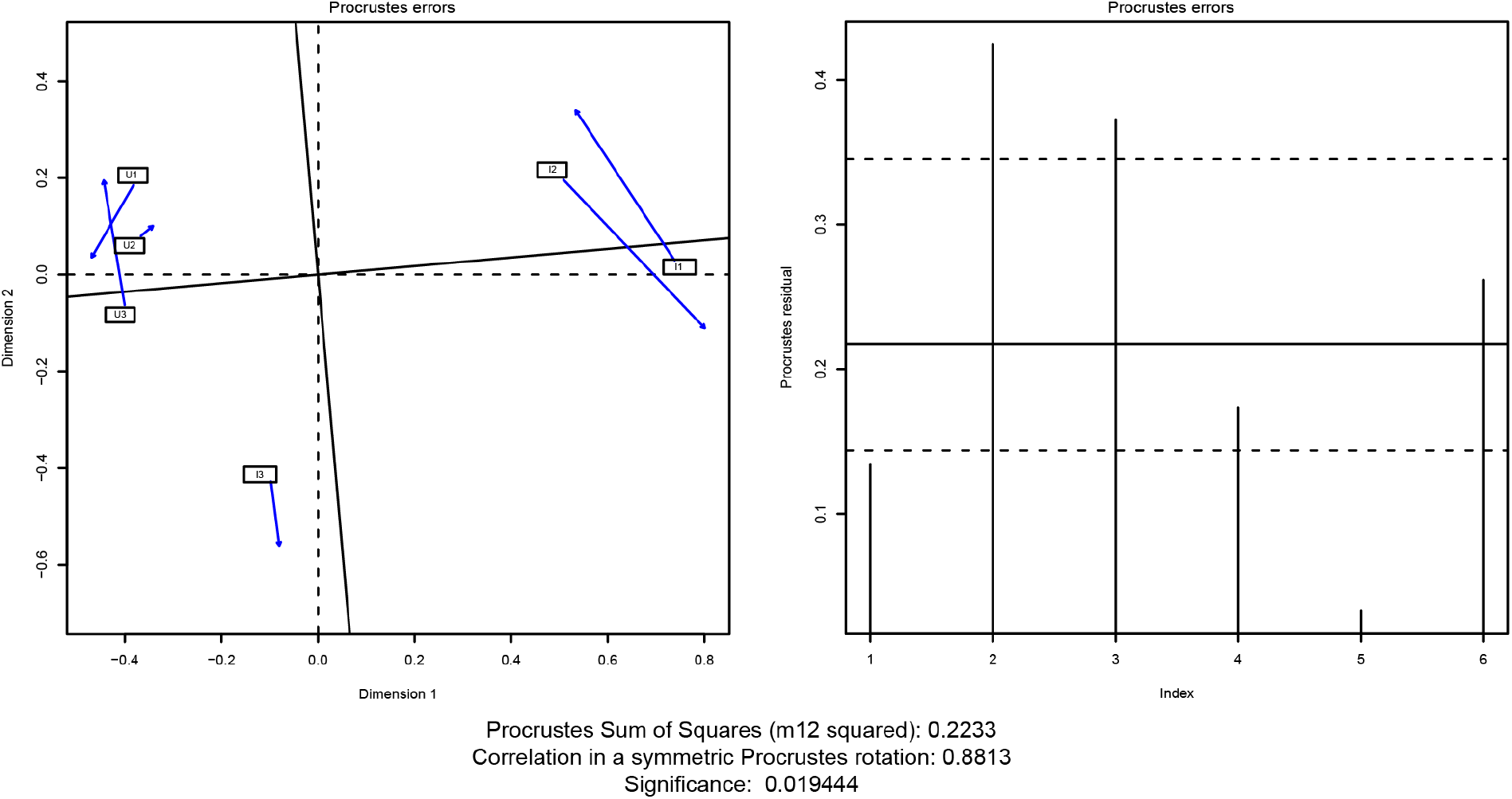
Procrustes analysis of dereplicated medium and high quality (dMQHQ) metagenome assembled genomes (MAGs) and viral metagenome assembled genomes (vMAGs). dMQHQ and vMAG relative abundance non-metric multidimensional scaling (NMDS) and viral genome-database NMDS showing high significant similarity of ordinations.

